# Early avoidance of human-dominated landscapes by juvenile Alpine golden eagles

**DOI:** 10.64898/2026.01.30.702719

**Authors:** Louise Faure, Yanni Gunnell, Hester Brønnvik, Enrico Bassi, Martin U. Grüebler, David Jenny, Petra Sumasgutner, Martin Wikelski, Kamran Safi, Elham Nourani

## Abstract

1. The expanding human footprint drives profound transformations in species, including not only morphological trait loss but also behavioural trait extinction. In contrast to behavioural trait extinction, which entails a reduction in genetic variability, behavioural plasticity maintains behavioural trait diversity and the associated genetic variability required for evolutionary adaptation. Yet, while behavioural plasticity shaping non-human capacity to coexist with humans has evolved under past, human-modified, environmental conditions, few studies have examined whether species retain a level of behavioural plasticity after past interactions with humans.
2. Using data from 64 juveniles golden eagles (*Aquila chrysaetos*) from a population demographically recovering after centuries of persecution, yet still restricted to less human-dominated alpine massifs, we tested whether human avoidance is innate or learned through experience during dispersal or parental influences during the pre-dispersal period. We performed a Principal Component Analysis (PCA) on landscape features used during the pre-dispersal and the first fifteen weeks of the dispersal phase, and used Mahalanobis distance as an indicator for quantifying changes in individual spatial behaviour over time.
3. The lower frequency of use of human-dominated landscape features, compared with the more sparsely populated mid- and high-elevation landscape features, resulted from individuals’ consistent use of similar landscape features across the pre-dispersal and dispersal periods. Avoidance of humans was therefore better explained by a learning or imprinting process prior to dispersal, which predisposed individuals to use landscape features similar to those experienced during pre-dispersal, rather than by learning during dispersal, or by an innate avoidance.
4. Imprinting on natal habitat features corroborates the observed limited range expansion of the alpine population beyond the massif borders. Differences among European populations in range recolonisation suggest contrasting histories of persecution, which may have left varying degrees of behavioural plasticity across populations.
5. As a guide for future conservation priorities, these findings underscore the need to reconstruct past conditions and/or to document contemporary animal behaviour in order to build a library of behaviours that can be used in the future to track behavioural changes.

## 1 Introduction

By modifying the adaptive landscape, humans have become the source of profound transformation in species (Dugatkin, 2024). Conspicuous morphological trait transformations have been drawing most of the attention. For example, African elephants (*Loxodonta africana*) in Gorongosa National Park, Mozambique, experienced a rapid loss of tusks following intensive ivory hunting (Campbell-Staton et al., 2021), while the enduring shortened wings in swallows have been considered an adaptation to collision risk associated with road traffic (Brown & Brown, 2013). In contrast and compared to morphological changes, behavioural traits are often assumed to be plastic and reversible (Wong & Candolin, 2015), and thus less prone to extinction or enduring transformation. However, changes in morphology can entail proximate and pleiotropic changes in behaviour, and vice versa. Selection for tameness in the silver fox, a melanistic variant of the red fox (*Vulpes vulpes*), for example, also selects for mottled coats, floppy ears, and curly tails (Dugatkin, 2018). Therefore, behavioural traits too can be exposed to enduring alteration or even extinction, a recent example of which is the loss of aggressive behaviour towards humans in the Apennine brown bear (*Ursus arctos marsicanus*) attributed to centuries of cohabitation with (and selection by) humans (Fabbri et al., 2025).

Extinction of certain traits increases individual vulnerability to future environmental change (Ghalambor et al., 2010), and can ultimately compromise species’ persistence. Since behavioural traits can have a genetic basis (Réale et al., 2007), a decline in the diversity of behavioural traits, like loss of morphological traits, could be associated with a reduction in genetic variability (Smith & Blumstein, 2013). A reduction that is likely to lengthen recovery times at population level, as more generations are required to rebuild genetic variability from a reduced gene pool, assuming that the trait and its genetic foundations have not been irreversibly lost (Loder, 2005;Conover et al., 2009). In contrast, behavioural plasticity—one form of phenotypic plasticity—refers to the ability of a single genotype to express different behaviours depending on environmental conditions without underlying genetic change (Hendry et al., 2008). By allowing organisms to adjust their phenotype during periods of rapid change, phenotypic plasticity may provide additional time for genetic adaptation to occur (Pigliucci, 2001; Tuomainen & Candolin, 2011) and may, under certain conditions, facilitate a population bouncing back to previous state.

Notable behavioural shifts have been attributed to behavioural plasticity, allowing species to use highly human-modified landscapes (Caspi et al., 2022). Contextual plasticity, defined as the immediate behavioural responses to environmental conditions, has enabled rural and urban silvereyes (*Zosterops lateralis*) to adjust their vocalisation frequency, amplitude, and duration when exposed to anthropogenic noise (Potvin & Mulder, 2013) and has also facilitated the innovation of foraging behaviours in blue tits (*Cyanistes caeruleus*) and great tits (*Parus major*), such as opening milk bottles to drink the cream (Lefebvre, 1995). Developmental plasticity, defined as experience-dependent changes that affect behavioural tendencies, has led white storks (*Ciconia ciconia*) in Europe to become less likely to migrate as they age, a phenomenon associated with the year-round availability of food at landfills (Andrade et al., 2025; Cheng et al., 2019). Similarly, transgenerational plasticity, where non-genetic parental influences shape offspring phenotypes, enables juveniles imprinted on the natal environment to use similar environments in which their chances of survival are higher (Davis & Stamps, 2004). For instance, red kites (*Milvus milvus*) from more artificial natal areas tend to capitalise on human resources available in urban environments during the prospecting phase (Orgeret et al., 2024). Conversely, a lack of behavioural plasticity can prevent species from associating to human-modified landscapes (Badyaev, 2005).

Contemporary human activities can influence the expression of behavioural plasticity by altering the environments in which individuals develop (Wong & Candolin, 2015). For instance, increased hunting pressure has led female elk (*Cervus elaphus*) to adopt movement strategies that minimise encounters with hunters (Thurfjell et al., 2017). In other words, the behavioural repertoire expressed by individuals today reflects past environmental conditions that favoured different responses (Buskirk, 2012; Sih et al., 2011). Yet, while several studies have emphasised the influence of past conditions in shaping the current behavioural repertoires of free-ranging populations (Tuomainen & Candolin, 2011), the role of past human pressures has been largely overlooked.

Following a period of systematic persecution in the 19th until mid-20th centuries (Bijleveld, 1974), the central European population of the golden eagle (*Aquila chrysaetos*) has recovered to the point of approaching carrying capacity (Fasce et al., 2011). Despite the recovery to population size saturation, their distribution is restricted to high mountain areas (Jenny et al., 2024), a pattern that is surprising given that golden eagles occupy diverse habitats from mountains to forests, meadows, and deserts – sometimes in close proximity to people (Bautista et al., 2024; Wentworth et al., 2024). Thus habitat type is arguably not the prime limiting factor in the expansion of the golden eagles’ distribution range. Instead, the limited range distribution in the central alps could suggest that this population specifically is not recolonizing the entirety of its ecological niche. Nourani *et al*. (2024) found that, although suitable areas for soaring flight for the Alpine eagle population increases with gains in flight experience, the realised distribution range however remains considerably smaller than predicted. The European Alps and their foreland areas have undergone increasingly intense environmental modification by human infrastructure (Gazzelloni, 2014), which raises the question of whether the restricted range of the eagles could be a result of avoiding humans. If correct, it remains unclear whether this avoidance behaviour, could be interpreted as a response to early exposure to current densities of humans in the region or as a legacy of past interactions with humans.

We used GPS technology to identify the landscape features used by 64 juvenile golden eagles during the pre-dispersal and over the first 15 weeks of the dispersal phase to understand whether (i) the population generally shows limited use of artificial landscapes from a young age, which could indicate avoidance behaviour selected at the population-level; (ii) there is a progressive reduction in the use of artificial landscapes during dispersal, which could indicate immediate learning of avoidance; and (iii) individuals use landscapes similar to the natal habitat, consistent with learning or imprinting prior to dispersal. If the hypothesis of a general avoidance of humans is supported (H1), an absence of variation in the use of landscape features between individuals is expected, whereas detectable variation in landscape features used would falsify it. If the hypothesis of immediate learning is supported (H2), within-individual temporal change, specifically a progressive reduction in the use of artificial landscapes during dispersal, is expected, whereas the absence of within-individual change would falsify it. If the hypothesis of learning prior to dispersal is supported (H3), an absence of differences between pre-dispersal and dispersal phases in landscape use is expected, whereas clear differences between phases would falsify it.

## 2 Materials and methods

### 2.1 Juvenile golden eagle tracking data

Sixty-four golden eagles were tagged at their nests in the Central Alps between 2017 and 2020 (see Supplementary Material, table S1). Tagging occurred between late June and mid-July, when juveniles were approximately 50 days old, as determined by plumage development (Bloom & Clark, 2001). Juveniles were equipped with Birds Solar UMTS tags produced by e-obs GmbH (Grünwald, Germany). The maximum combined mass of the leg-loop harness and logger was 60 g, i.e., less than the recommended 3% of the bird’s body weight at the time of tagging (Kenward, 2000). The loggers collected one GPS point every 20 minutes (Zimmermann et al., 2025). The tags had a low median horizontal error of 12.19 m and a median dilution of precision (DOP) (a measure of the effect of satellite geometry on the accuracy of GPS positioning) of 4.39 m during the pre-dispersal period and the first 15 weeks of dispersal, corresponding to a ‘good’ level of location accuracy (Isik et al., 2020).

Tracking data are part of a life tracking study. The analysis focused on two periods in the eagles’ life history: (i) the pre-dispersal period, during which juveniles remain dependent on their parents, which starts with fledging (i.e. leaving the nest) and ends with emigration from natal territory, and (ii) the subsequent dispersal period, which starts with emigration from the natal territory and is restricted in this study to the first 15 weeks during which juveniles’ movements are less constrained by avoidance of territorial adults that are still in parental care mode (Watson, 2010). The timing of fledging was estimated from changes in first-passage time (FPT) (Fauchald & Tveraa, 2003). FPT is the time it takes an animal to move through an area of a given radius, for example an animal moving in a rapid, directed manner has lower FPT than one moving slowly and in an undirected manner. Here, fledging was defined as the first change from high to low FPT on the assumption that eagles will start moving through a given radius only after fledging. To allow for differences between individuals, FPT was calculated for each individual in two steps using the *adehabitatLT* R package (Calenge, 2006). First, FPT was calculated at radii between 20 m and 2020 m at 50 m increments and the radius of the first inflection in the log variance in FPT over space was retained. This meant that each individual could have its own radius based on the data it sent. Second, smooth splines using 30 knots were fitted to FPT at the retained radius over time. The first minimum after a maximum in the smooth spline was retained as the fledging date. Emigration dates were based on Nourani *et al*. (2024).

### 2.2 Terrain-based features of the landscape

To identify the landscape features the golden eagles used, a series of terrain-based features was calculated. Landscape units were established according to subdivisions recommended by the Living Landscape Project (Warnock & Griffiths, 2015), and simplified into three categories: landform, land cover, and settlement pattern. Within each of these three categories, terrain-based variables were selected according to previously identified determinants of space use among golden eagles (topographic heterogeneity as a proxy for orographic updraughts generated by the deflection of air currents by a physical barrier (Bohrer et al., 2012; Murgatroyd et al., 2018), vegetation cover type (Singh et al., 2016; Whitfield et al., 2007), and human-made infrastructures (Nielson et al., 2016; Sergio et al., 2006; Tack et al., 2020). Topographic, land cover and settlement pattern categories were described by 18 variables derived from a range of data-sets summarised in Table 1. All the variables are quantitative. Categorical variables such as landcover types, were converted to continuous values by determining proportion (in %) of GPS location points within each land cover type during each individual flight sequence.

**Table 1.**
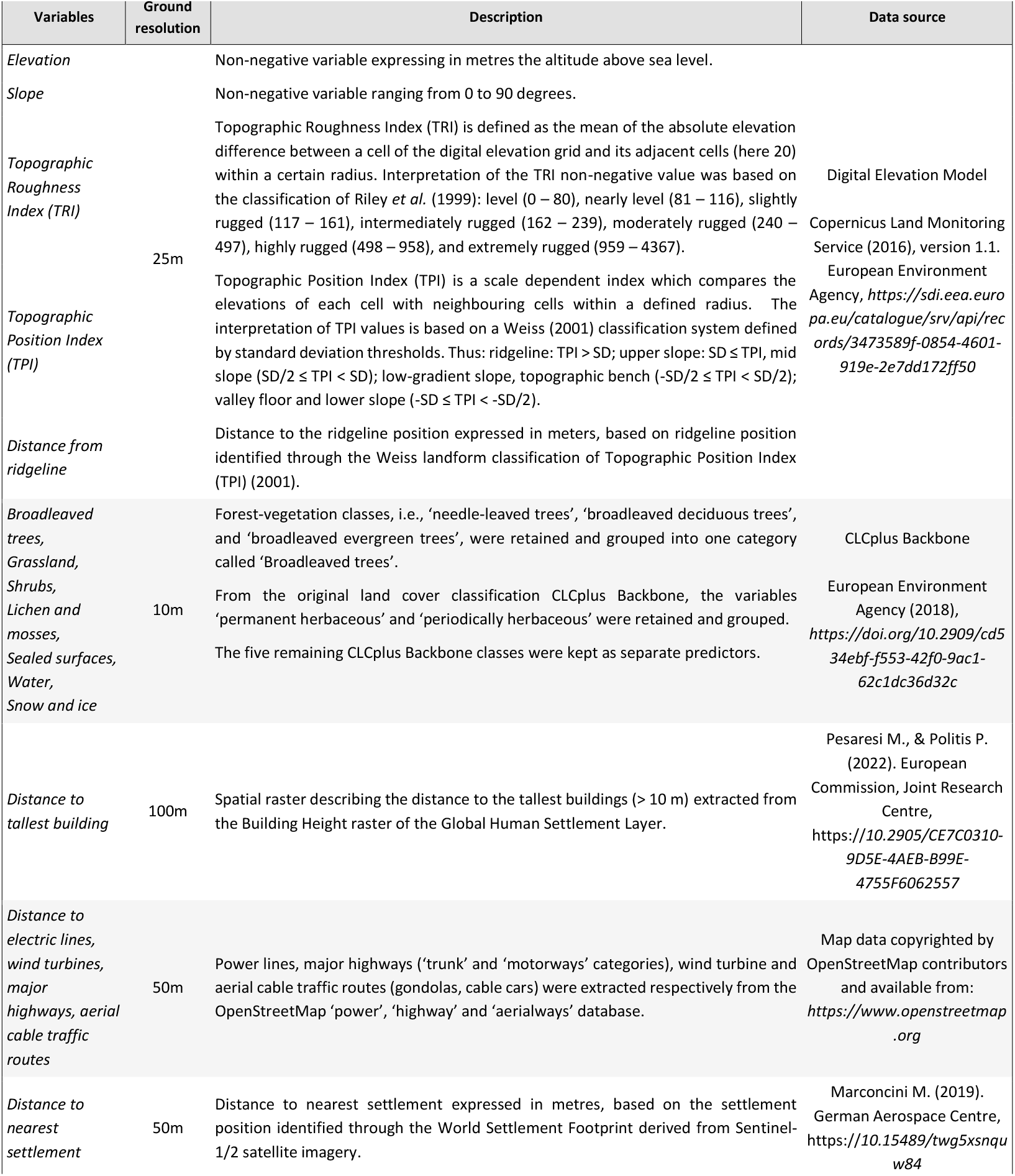
Topographic and landform variables for quantifying the landscape categories frequented by young golden eagles. Variables derived from the 25 m resolution European Digital Elevation Model (EU-DEM), version 1.1, 2011, European Environment Agency (EEA)

Topographic variables, i.e., elevation, slope, Topographic Position Index (TPI; Weiss, 2001), Topographic Roughness Index (TRI, Riley et al., 1999), distance to ridge-line, were extracted from a 25-m Digital Elevation Model of the Alps (Copernicus, https://land.copernicus.eu/) using SAGA-GIS. Ridge-lines were delineated from multi-scale TPI where the TPI with an inner radius of 10 m and an outer radius of 200 m was retained. Distances to ridge-lines were computed with SAGA’s ‘proximity raster’ function (Conrad et al., 2015) and were then used to attribute to each of the raster cells a distance to the closest ridge-line. Distances to human-built infrastructures were derived through a similar approach: the 10 m World Settlement Footprint (Center, 2021), clipped to the extent of the Alpine study area, was aggregated to 50 m cells, and OpenStreetMap vector features (OpenStreetMap contributors, 2025) were contoured by a 50 m buffer, and converted to distance values.

### 2.3 Measuring variation between individuals: identification of various landscape utilisation strategies

#### 2.3.1 Track sampling and individual weights

The locations of the juvenile golden eagles were filtered on the basis of three criteria relating to (i) timing of dispersal (see Juvenile golden eagle tracking data), (ii) time lapse between locations, and (iii) minimum distance between two locations. GPS data corresponding to the dispersal period were split into 7-day sequences. Within each week-long sequence, eagle location points were selected based on two conditions: (i) they should be least 15 minutes apart (consistent with the original 20-minute measurement interval, thus removing any high-resolution GPS bursts); and, in accordance with the coarser pixel resolution of the raster layers, (ii) they should be separated by a minimum distance of 150 m. Individuals with at least 30 location records per week were retained. To account for variability in data collection among sampled individuals, a weighting index based on the percentage of locations relative to the total number of locations for each individual was generated.

#### 2.3.2 Principal Component Analysis

A Principal Component Analysis (PCA) was performed in order to identify groups of correlated variables and to highlight differences in spatial behaviour among the juvenile golden eagles (Abrahms et al., 2017; McGarigal et al., 2000). The first step involved extracting landform, land cover, and settlement raster values at each individual’s location. Terrain characteristics were then summarised for each individual eagle by calculating the median, maximum, minimum, variance, and first quartile of the collected values. To facilitate comparisons between variables expressed in different measurement units, each variable was standardised by dividing it by its standard deviation. Variables were then filtered through a three-step procedure that balanced biological interpretability with statistical redundancy and importance to the PCA components. For each variable, correlation matrices among its set of metrics (median, minimum, maximum, variance, first quartile) were generated, and redundant metrics were removed. A Pearson correlation matrix which measures the linear dependence between two variables, was then computed across the remaining variables to eliminate highly correlated pairs (defined here when |*r*| > 0.60 ± 0.03) (Kassambara, 2017b) (see Supplementary Material, fig. S1). Finally, variables with low contribution, i.e., low contribution to the principal components and low representation quality, were identified using the weighted contribution and squared cosine metrics (Abdi & Williams, 2010). Variables with low values on the leading components were considered for removal (Faure, 2025b).

Entering the resulting dataset into the PCA using the FactoMineR package (Lê et al., 2008), eigenvalues were extracted to determine the number of principal components. This was achieved by identifying the point beyond which the remaining eigenvalues became relatively small and of comparable size (Jolliffe, 2002; Peres-Neto et al., 2005), thereby further allowing a reduction in dataset dimensionality without losing significant information. Based on the selected principal components, individual points summarising a 7-day sequence were then grouped into clusters using a *k*-means algorithm, a centroid-based clustering method that partitions an individual eagle’s position into *k* clusters by minimising the distance between each point and the centre of its nearest cluster. Landscape Utilisation Strategies (LUSs), defined as the consistent occurrence of individuals within a 7-day sequence amid certain landscape features, were identified on the basis of clusters, each representing a combination of landscape features used. The number of strategies was determined on the basis of (i) cluster strength indicators (Kassambara, 2017a) and (ii) interpretability of the resulting strategies from a biological perspective. Statistical strength indicators included within-cluster distances, distances between cluster centres, multi-index cluster validation using the NbClust R package (Charrad et al., 2014; Kassambara, 2017a), and the Bayesian Information Criterion (BIC; Schwarz, 1978).

### 2.4 Measuring plasticity in landscape use

#### 2.4.1 Metrics of individual behavioural change

Change in behaviour was measured using the Euclidean distances between points in the principal-component space, not accounting for week-to-week geographic distances and available landscape features. By retaining a larger number of uncorrelated principal components, these Euclidean distances approximate Mahalanobis distances in the original multivariate space, which account for the correlation coefficient between variables (Mahalanobis, 1936). For each individual, distances were computed between successive 7-day positions during the first 15 weeks of dispersal, and between the pre-dispersal position and each 7-day position of the first 15 weeks of the dispersal period. Distances were summarised for each individual by the mean, yielding one value for week-to-week change and one for pre-dispersal to dispersal change.

#### 2.4.2 Testing for individual plasticity through accumulation of experience during dispersal

Our hypotheses posited a change in behaviour with experience, whereas the null hypothesis stated that golden eagles do not learn to avoid human-dominated areas, as would be indicated by behaviour that remains consistent over time. To test whether individual behaviour is more or less consistent than expected by chance, a random dataset was created. Principal-component coordinates were shuffled among individuals within the same dispersal stage, accounting for ontogenetic change (Nourani et al., 2024). The Mahalanobis distances were then calculated and summarised for each individual by the mean. Randomisation was repeated 1,000 times. Statistical significance was assessed as the proportion of random Mahalanobis distances that were smaller or equal to the observed Mahalanobis distances, and results were considered significant at 0.05. Analyses were restricted to the 47 individuals with data covering at least two weeks of the dispersal period.

#### 2.4.3 Testing for pre-dispersal behaviour as a determinant of golden eagle behaviour during dispersal

Our hypothesis was that dispersal behaviour remains similar to pre-dispersal behaviour, with the null hypothesis stating no such similarity. Accordingly, observed distances from each dispersal week to the individual’s pre-dispersal position were expected to be shorter than distances obtained under a null distribution. A second null distribution was built by shuffling principal-component coordinates among different individuals within the same dispersal stage, and recomputing the Mahalanobis distance to each individual’s own pre-dispersal position. This procedure was repeated 1,000 times and distances were summarised by the mean, producing one value for each individual and for each randomisation. Statistical significance was assessed as the proportion of Mahalanobis distances that were smaller or equal to the randomly obtained Mahalanobis distances, and results were considered significant at 0.05. Analyses were restricted to the 49 individuals with data covering the pre-dispersal period and at least one week of the dispersal period.

## 3 Results

### 3.1 Individual variations in landscape utilisation strategies

While considering only the first four principal components, which together explained 70.6% of the variance of the dataset (see Supplementary material, fig. S2), three landscape utilisation strategies based on 12 variables arose from the statistical clustering process (see fig. 1). The first two components explained 28.1% and 20.2%, respectively, with the first capturing behavioural variations linked to the presence of human-built infrastructures such as roads, wind turbines, electric lines and aerial cable traffic routes. The second component described a gradient from sparsely to highly vegetated areas (see Supplementary Material, fig. S3).

**Figure 1.**
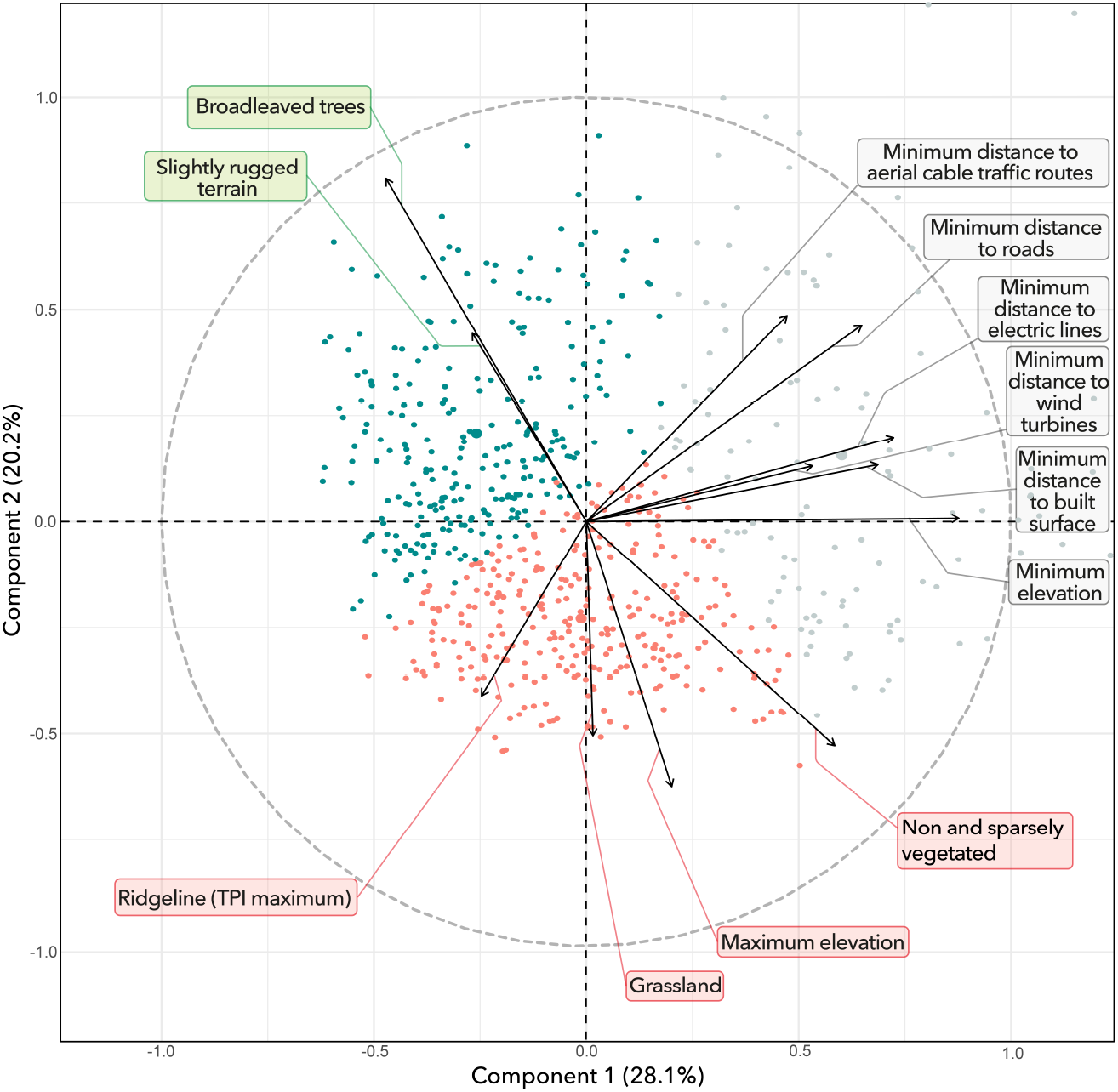
PCA loadings plot. The graph shows the relationship between the five principal components and the original variables. Each point represents one week of an individual’s life during its dispersal and pre-dispersal periods (not distinguished in this graph). Red squares, green triangles and grey circle represent the Rangetop, Forest and Valley strategies, respectively. Arrows point to various degrees in the direction of, or away from, components 1 and 2. Arrow lengths indicate the contribution of the original variable to the principal component. The factor circle (dashed lines) represents the boundary of maximum contribution of the original variables to components 1 and 2.

The most frequent landscape utilisation strategy (313 weeks in the PCA plot, Figure 1), hereafter labelled as the Rangetop Strategy, was defined by three positively correlated variables: ‘Maximum distance to ridgelines’, ‘Maximum elevation’, and ‘Absence of vegetation cover’. Birds displaying a Rangetop Strategy used the elevation belts where woody vegetation is scarce, i.e., alpine meadows above the treeline.

With a total of 258 weeks, the use of low-elevation and forested mountain terrain with a low human footprint was the second most frequent landscape utilisation strategy. Here referred to as the Forest Strategy, this pattern was defined as a combination of two highly correlated variables—slightly rugged terrain (‘Mean Topographic Roughness Index’) and ‘Broadleaved trees’. This group of eagles was conspicuously confined to the negative region of the PCA loadings plot relative to the location of the other two landscape utilisation patterns (Figure 1), thus highlighting an affinity with the lower montane vegetation belts suited to hosting deciduous trees or at least mixed stands of hardwood and softwood species.

With only 118 weeks in the PCA, the third and least frequent landscape utilisation strategy highlighted cases of eagle occurrence at the smallest recorded distance to human infrastructures. This was defined as a Valley Strategy because the valley floors and lower slopes of the central Alps concentrate the highest densities of human construction, activity, and infrastructure (Figure 1).

The three clusters were validated by a combination of strength indicators and interpretability. Distance indicators suggested an optimal number of clusters of less than five. Beyond this threshold, the distance between points within the same cluster increased while the distance between clusters decreased, thereby leading to clusters with unclear borders and an increased risk of point misclassification. The Bayesian Information Criterion (BIC) suggested five, three or six optimal clusters (see Supplementary Material, fig. S4). We opted for the three-cluster partitioning solution because the five and six cluster variants were more difficult to interpret, with little additional gain in explaining the total variance.

### 3.2 Individual consistency in landscape use across the pre-dispersal and first 15 weeks of the dispersal period

Avoidance of human-dominated areas is best explained by natal conditions experienced before dispersal. Individuals exhibited consistency in landscape use during the first 15 weeks of dispersal. Overall, 29 individuals were found to have specialised in one predominant strategy during dispersal, whereas 5 individuals adopted all three strategies in relatively equal proportions (Figure 2). Likewise, among the 34 individuals with complete data transmission, 28 showed significant consistency (*p* ≤ 0.05) in landscape use during the first 15 weeks of dispersal, therefore not supporting a learning-driven avoidance of human-dominated areas during dispersal. Moreover, pre-dispersal behaviour predicted dispersal behaviour, i.e., the landscape characteristics used during dispersal were similar to those experienced during pre-dispersal. Among the 34 individuals with complete records for both the pre-dispersal period and the subsequent 15 weeks of dispersal, 16 individuals displayed significant consistency in their use of the landscape features across the pre-dispersal and the consecutive 15 weeks of the dispersal period (*p* ≤ 0.05, Figure 2). Details of the individual *p*-values are available in the Supplementary material (Figure S4; Table S2).

**Figure 2.**
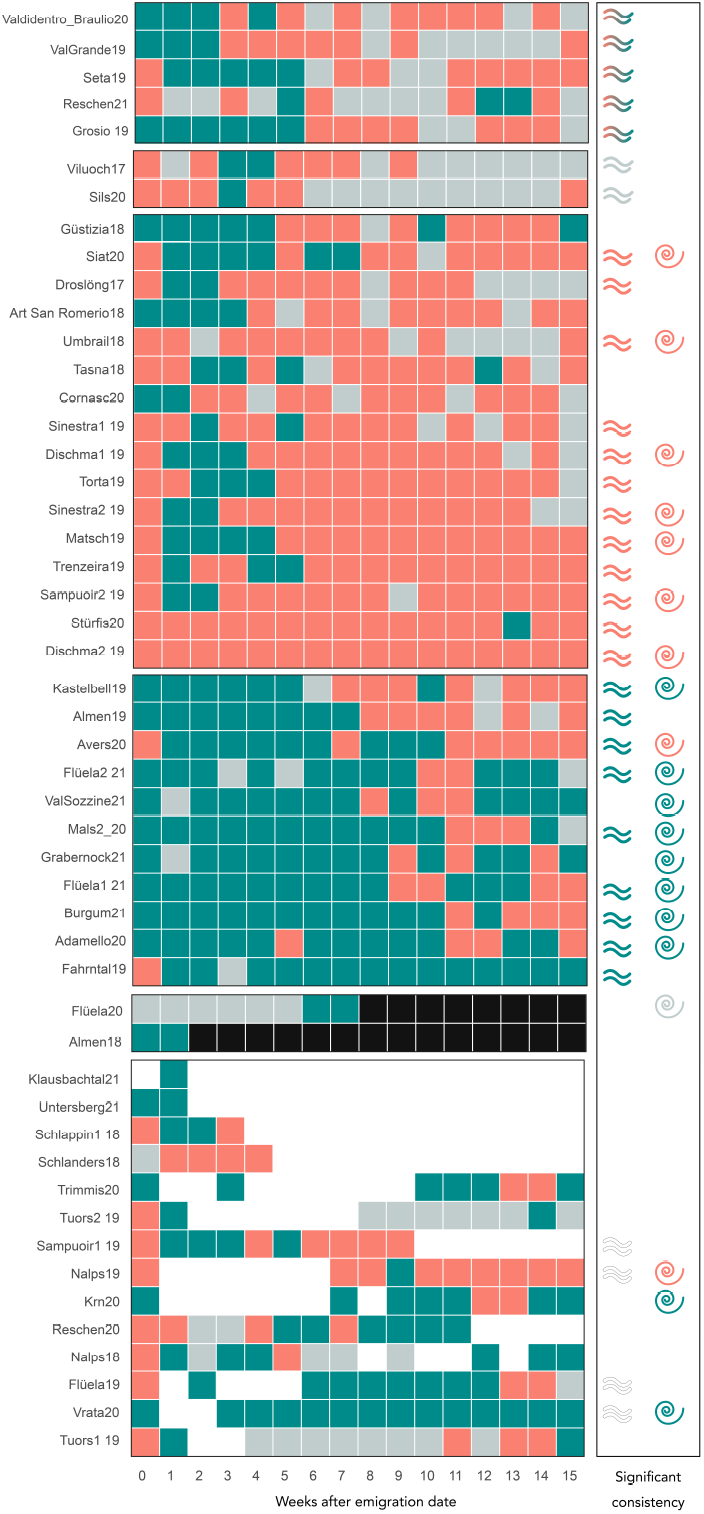
Consistency in landscape utilisation strategy (LUS) during the pre-dispersal and the first fifteen weeks of the dispersal period. Left column lists the individual eagles. Red, green, and grey represent Rangetop, Forest, and Valley strategies, respectively. Black corresponds to deceased birds and white indicates gaps in data transmission. Week 0 defines the maintained phase of the pre-dispersal period. Individuals have been grouped according to their predominant strategy during dispersal and incomplete data transmission due to logger related issues or identified death. The equal sign and spiral symbol indicate significant consistency (*p* ≤0.05) in landscape utilisation strategy from one week to the next during the dispersal phase, and between the pre-dispersal period and each week of the dispersal phase, respectively.

## 4 Discussion

A dataset of 64 juvenile golden eagles was analysed to assess how selectively individuals responded to natural and human landmarks during the transient phase of dispersal, a time during which juveniles encounter previously unknown landscape features. Results indicated that, among the three identified landscape utilisation strategies, the Valley strategy, corresponding to landscape features dominated by human presence and activity, was used least frequently. Limited use of artificial environments was best explained by pre-dispersal learning (or imprinting), which predisposed individuals to use familiar landscape features, rather than by learning during dispersal or by an innate avoidance.

The influence of early exposure to particular landscape features on later used landscape features suggests that golden eagles are subject to a process of natal habitat imprinting (Davis & Stamps, 2004). Imprinting on natal habitat features is consistent with observations of Alpine golden eagles often returning to within a 120 km radius of their natal territories to establish their own nesting places (Haller, 1996). Both the location of the nest site in areas of low disturbance (Jenny et al., 2024) and parental spatial behaviour, since parents are reported to fly and guide their offspring to carrion (Hemery et al., 2023; Ricau & Decorde, 2009), might have reduced juveniles’ exposure to artificial landscape, resulting in a limited use of the Valley strategy later in life.

Natal habitat imprinting may explain, and be confirmed by, the limited expansion of our study population beyond the Alps (Fasce et al., 2011). The consequences of this inertia in range expansion include increased densities of golden eagles and thus increased competition between individuals, which in turn lead to reduced fecundity (Chambert et al., 2020). Increased densities of golden eagles also drive increased predation pressure on prey and mesopredators, potentially affecting herbivore-mesopredator interactions, as observed in the Finnish population (Lyly et al., 2015). Restricted range expansion is, however, not a common feature amongst golden eagle populations since, for instance, the similarly hunted population of the Iberian Peninsula is expanding (Martínez-Abraín et al., 2019). Iberian golden eagles also shifted to tree nest sites, interpreted as a release from past human pressure (Martínez-Abraín et al., 2021). Population differences in range expansion under relaxed human pressure call for re-evaluating the causes of these variations and their underlying factors.

Variation in range expansion across populations could result from differences in behavioural plasticity and from environmental conditions encountered in the past. First, past conditions, particularly hunting pressure in the Alps, may have favoured the inter-generational transmission of human-averse traits from experienced survivors over more costly individual learning (Donelan et al., 2020; Harmon & Pfennig, 2021; Kuijper & Hoyle, 2015). Second, limited behavioural plasticity may stem from differences in the population’s genetic make-up, consistent with the reduced genetic variability observed in Mediterranean lineages compared with Holarctic golden eagle lineages (Karabanina et al., 2024, but see Nebel et al., 2019). This reduction in genetic variability could have been selected by past changes in survival rates or in the fitness of particular traits (Tuomainen & Candolin, 2011). For instance, high prey availability at high elevations (Sergio et al., 2006) in open habitats maintained by historical pastoralism (Pedrini & Sergio, 2001) may have indirectly increased the fitness of the Rangetop strategy and thereby promoted its transmission across generations. Conversely, reduced survival in the vicinity of humans could also have directly selected for human-averse traits, as proposed for the Iberian Peninsula population, which persisted in inaccessible, sparsely populated territories through periods of intense persecution (Fernández-Gil et al., 2023).

## Conclusion

Using data from a recovering population of golden eagles, which, after intensive hunting, has increased demographically but not expanded its range, this study tested whether this limited range expansion could be explained by avoidance of humans. Results show that the juveniles displayed a limited use of human-dominated landscape features, as indicated by the low cumulative rates of valley use (Valley strategy) compared with the more sparsely populated mid- and high-elevation habitat mosaics (Forest and Rangetop strategies). Consistent use of landscape features similar between the pre-dispersal and dispersal phase, further resulting in avoidance of human-built landscape, suggest a learning process occurring during pre-dispersal rather than through accumulation of experience during the post-dispersal phase. The influence of early experience on later space use, also known as the natal habitat imprinting hypothesis, corroborates the observed limited range expansion of the alpine population beyond the massif borders. Variations of range expansion between populations of golden eagles may result from differences in behavioural plasticity, which has evolved under various historical context of human animal interactions. Because behavioural plasticity influences a species’ capacity to persist under future environmental change while being determined by past conditions, reconstructing past conditions or, alternatively, documenting contemporary animal behaviour to which future observations can be compared is essential.

## Author Contribution

Elham Nourani, Louise Faure, and Kamran Safi conceived the study. Yanni Gunnell provided supervision, insight from a geographical perspective, and manuscript editing. Kamran Safi, Martin Wikelski, David Jenny, Martin U. Grüebler, Enrico Bassi and Petra Sumasgutner contributed the data. Hester Brønnvik and Louise Faure processed the GPS data. Louise Faure processed the terrain-based data, carried out the data analysis, and wrote the first draft of the manuscript. All authors discussed the findings and commented on the structure and wording of the manuscript.

## Acknowledgements

Louise Faure and Elham Nourani were supported by the Deutsche Forschungsgemeinschaft (DFG) under Germany’s Excellence Strategy (– EXC 2117 – 422037984). Louise Faure also received support from the Graduate Initiative “Biodiversité et Bioressources” at Claude Bernard University Lyon 1, funded through France 2030 by the Agence Nationale de la Recherche (ANR), reference “ANR-21-SFRI-0001”. Elham Nourani was supported by the PRIME program of the German Academic Exchange Service (DAAD) with funds from the German Federal Ministry of Education and Research (BMBF) and the Swiss National Science Foundation (Starting Grant TMSGI3_226462). We thank Ross Purves for supporting and facilitating this interdisciplinary collaboration, as well as for offering his fresh perspective on the project. We acknowledge the institutions and persons who made fieldwork and data collection possible — as listed in the acknowledgements of Nourani et al., 2024. We are grateful to the Walter & Bertha Gerber Foundation and Yvonne Jacob Foundation for financially supporting the research of David Jenny and Martin U. Grüebler.

## Data availability statement

The data needed to replicate the findings of this paper have been uploaded on the Edmond repository (Faure, 2025a). A DOI will be generated and shared pending the acceptance of the paper for publication. The code for data analysis is accessible via the github repository ‘Landscape use golden eagles’ (Faure, 2025b).

## Ethics

The capture and ringing of golden eagle juveniles were carried out in accordance with national and regional regulations in each country. In Austria, all procedures were reviewed and approved by the Ethics Committee of the University of Vienna (No. 2020-008) and authorised by the Federal Ministry for Education, Science and Research (No. 2020-0.547.571), as well as by regional authorities in Styria (BHLI-165942/2021-2) and Upper Austria (LFW-2021-263262/7-Sr). In Italy, permissions for handling and tagging were granted by the Autonomous Province of South Tyrol (Decrees 12257/2018 and 8788/2020) and by the Region of Lombardy, with additional approval from the Istituto Superiore per la Protezione e la Ricerca Ambientale under the “Richiesta di autorizzazione alla cattura di fauna selvatica per scopi scientifici” (l.r. 26/93). In Switzerland, activities were authorised by the Federal Food Safety and Veterinary Office Grisons, permit no. GR 2017_06, GR 2018_05E, GR 2019_03E, GR/08/2021, and the Federal Office for the Environment. In Germany, permits were issued by the government of Upper Bavaria (2532.Vet_02-16-88 and 2532.Vet_02-20-86) for bird handling, tagging, and ringing. All handling procedures adhered to the ASAB/ABS guidelines for the ethical treatment of animals in behavioural research, as well as to all relevant institutional, national and international regulations on animal welfare. The handling of golden eagles was performed with the utmost care and with and minimal disturbance to nests and the surrounding landscape. Ethical approval for the inclusion of animals in this research was obtained from the scientific commission of the Swiss Ornitho-logical Institute and the respective national authorities, and complied with European Commission and Federation of European Laboratory Animal Science Associations (FELASA) guidelines.

## Supplementary material

**Figure S1.**
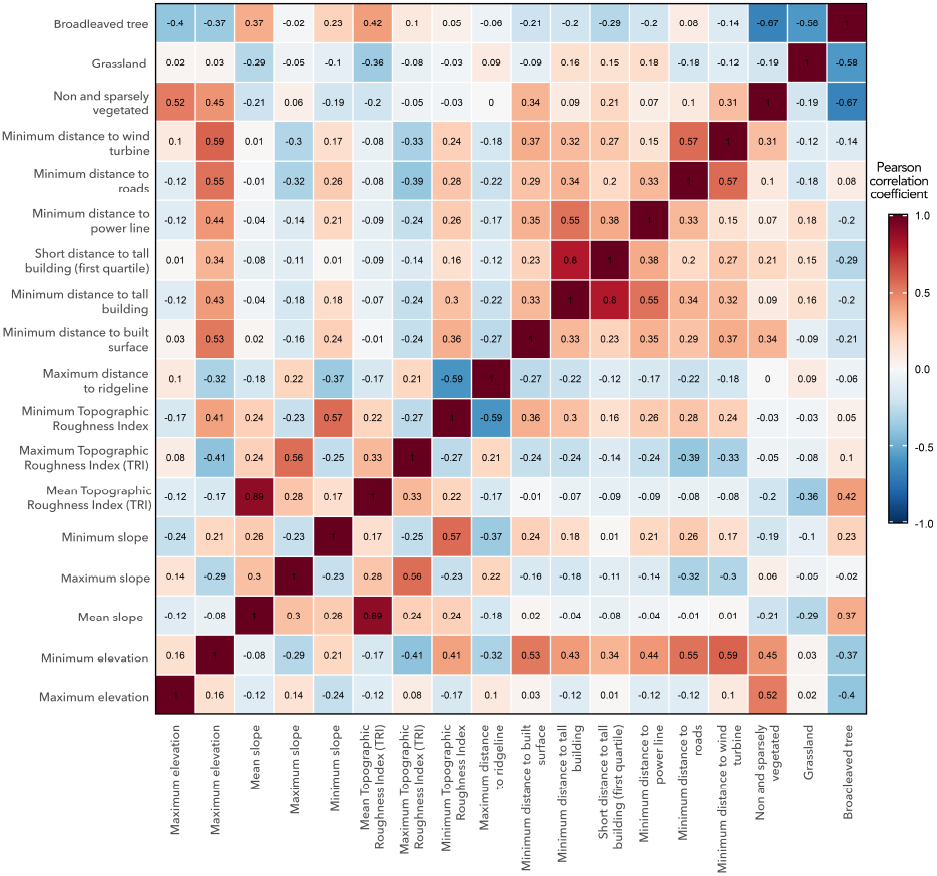
Correlation matrix between variables selected for this study. The display provides and overview of the variables and correlation strengths between them using the Pearson correlation coefficient.

**Figure S2.**
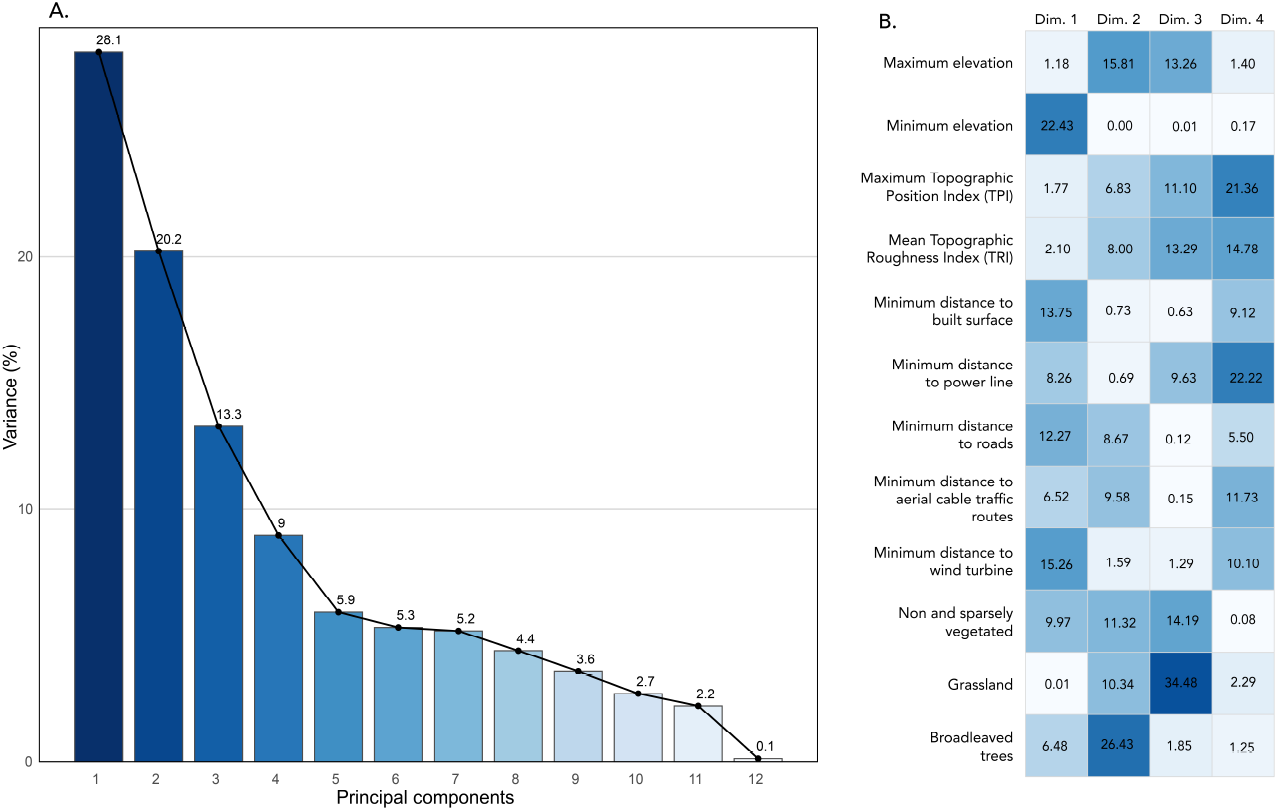
Eigenvalues and relative contribution to the first four components of the PCA. A: scree-plot of the eigenvalues relative to the 12 variables; B: relative contribution of different variables ex-pressed in percentage of the four components of the PCA. Figures highlight the variables with the greatest influence on shaping a particular principal component. The first two dimensions (Dim.) explain 48.3% of the total variability observed in the dataset.

**Figure S3.**
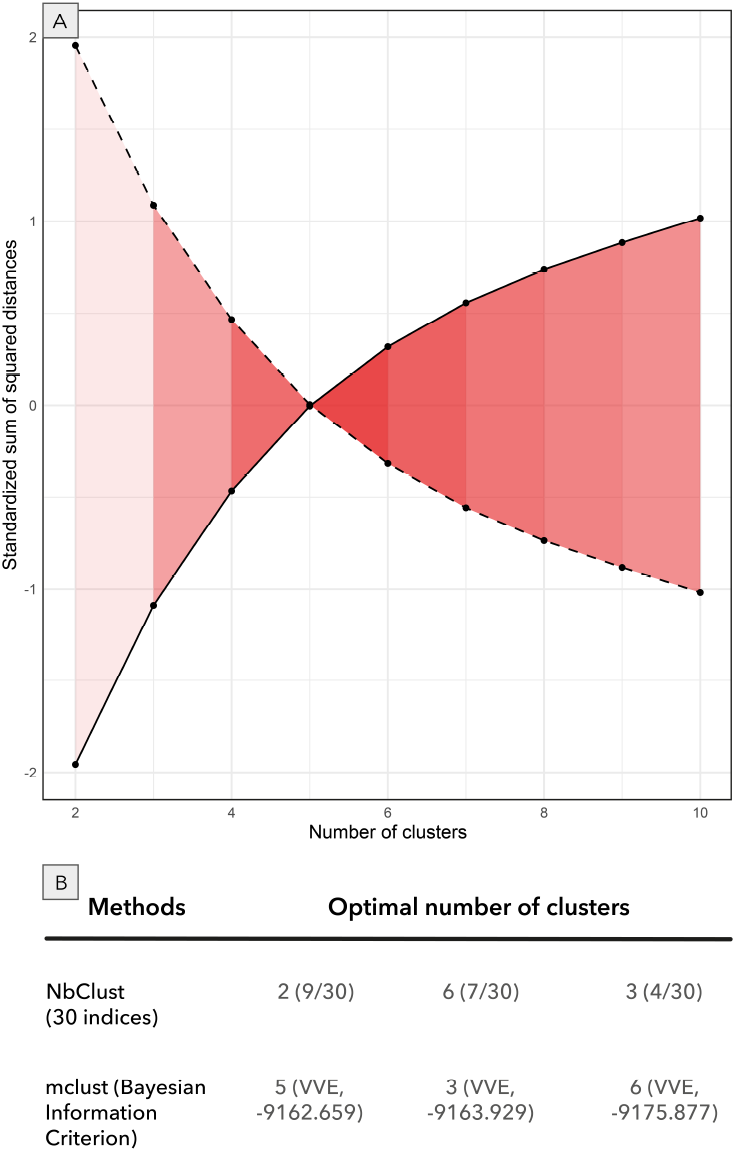
Cluster strength indicators. A: Optimal number of clusters based on distance within and between clusters. Solid line represents the sum of squared distances between clusters. Dashed line represents the sum of squared distances within clusters. Intensity of the red colour increases with the suitability of the corresponding number of clusters. B: Determination of the optimal number of cluster.

**Figure S4.**
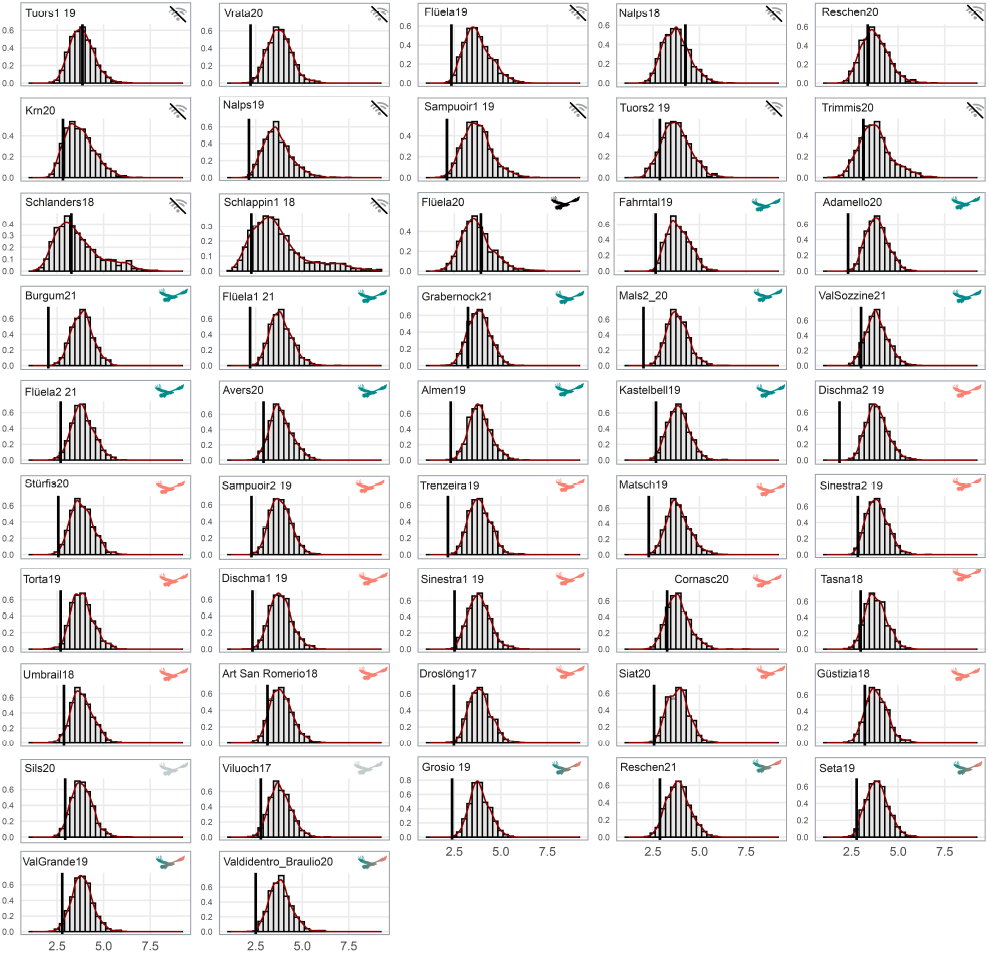
Null and observed distributions of the mean of the distance between consecutive weeks of the dispersal phase. Null distributions of mean Mahalanobis distances between consecutive 7-day sequence point during dispersal (x-axis), based on 1000 within-week random permutations. Kernel density estimates represented by a red-curve and histograms represent the permuted values (y-axis). The vertical black line marks the observed mean distance for each individual. Eagle icon colours reflect predominant landscape-use strategies: green for forest specialists, red for rangetop specialists, and grey for valley specialists. Transmission symbol denotes individuals that ceased transmission or lost their logger; black icon indicates mortality; multicolour icons denote generalists.

**Table S1.**
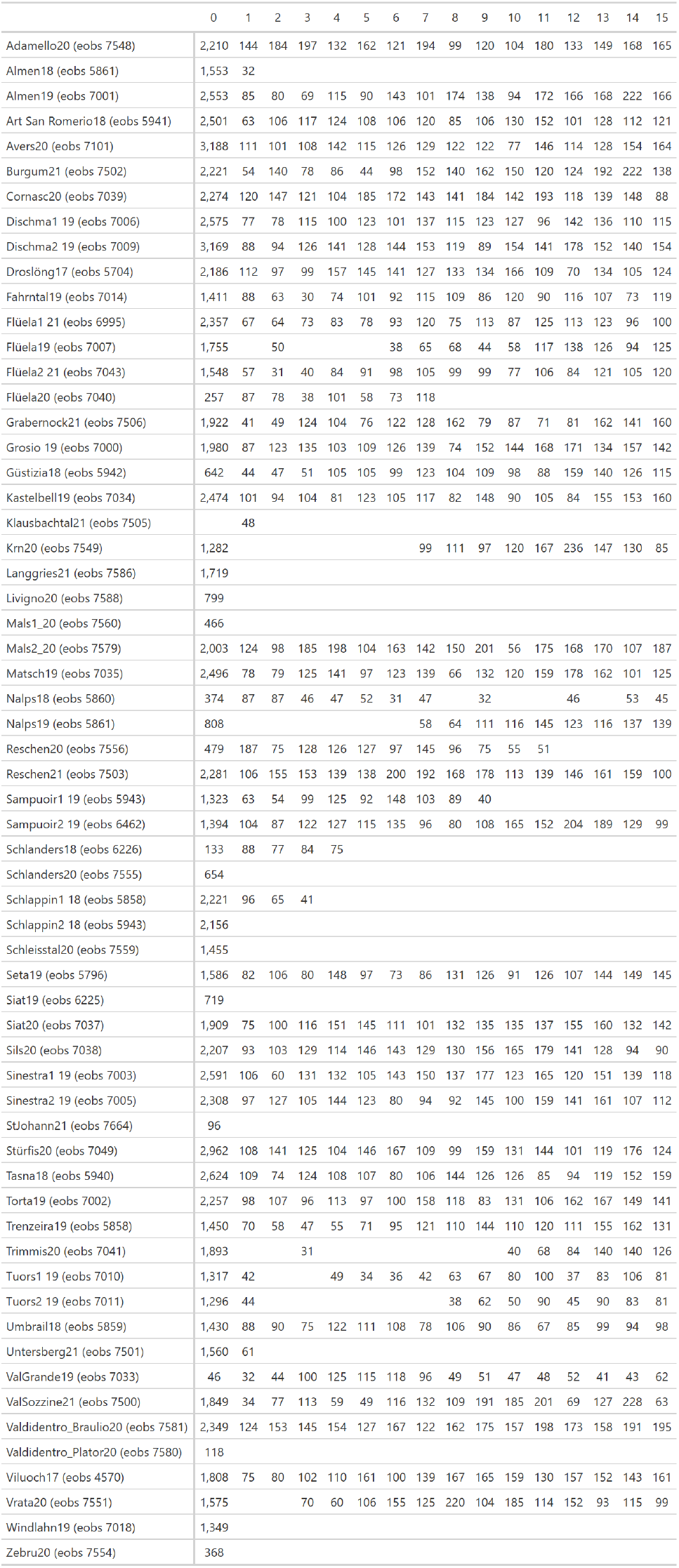
Number of GPS locations per individual per weeks.

**Table S2.**
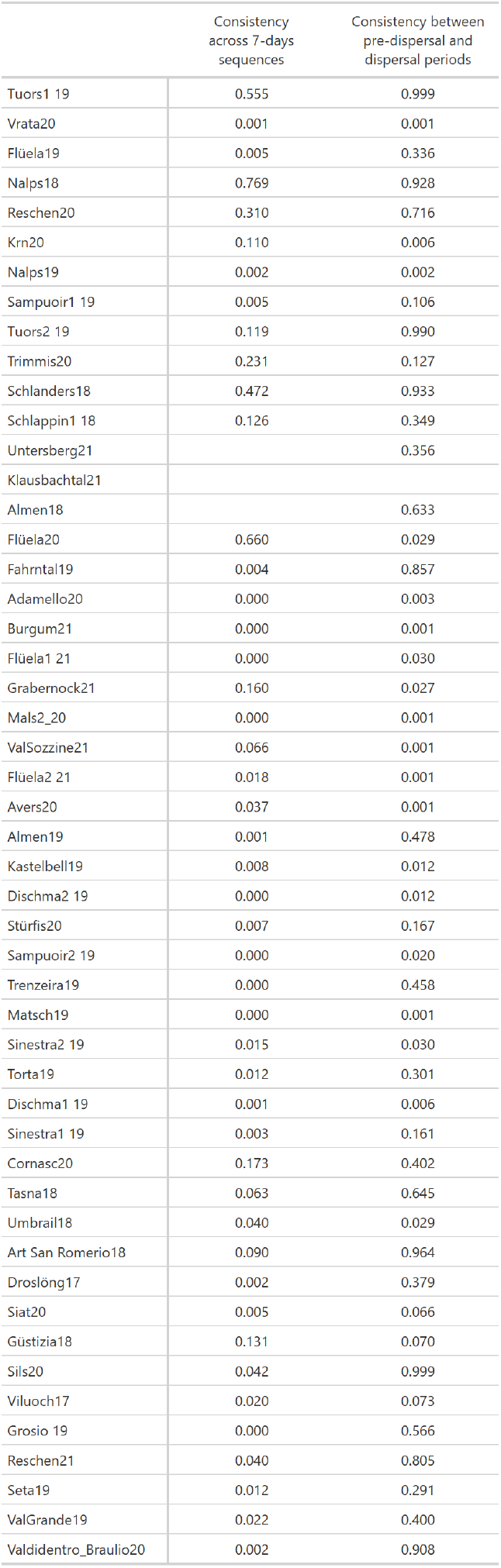
Inventory of *p*-values per individual.

## References

Abdi, H., & Williams, L. J. (2010). Principal component analysis. Wiley Interdisciplinary Reviews: Computational Statistics, 2 (4), 433–459. 10.1002/wics.101

Abrahms, B., Seidel, D. P., Dougherty, E., Hazen, E. L., Bograd, S. J., Wilson, A. M., Weldon McNutt, J., Costa, D. P., Blake, S., Brashares, J. S., & Getz, W. M. (2017). Suite of simple metrics reveals common movement syndromes across vertebrate taxa. Movement Ecology, 5 (1), 12. 10.1186/s40462-017-0104-2

Andrade, P., Franco, A. M. A., Acácio, M., Afonso, S., Marques, C. I., Moreira, F., Carneiro, M., & Catry, I. (2025). Mechanisms underlying the loss of migratory behaviour in a long-lived bird. Journal of Animal Ecology, 95 (5), 1061–1075. 10.1111/1365-2656.70035

Badyaev, A. V. (2005). Stress-induced variation in evolution: From behavioural plasticity to genetic assimilation. Proceedings of the Royal Society B: Biological Sciences, 272 (1566), 877–886. 10.1098/rspb.2004.3045

Bautista, J., Gómez, G. J., Otero, M., & Garrido, J. R. (2024). Golden eagle in souhtern spain recolonize human-dominated landscapes. In D. H. Ellis, J. Bautista & C. H. Ellis (Eds.), The golden eagle around the world: A monograph on a holarctic raptor (pp. 221–252). Hancock House Publishers, Surrey (BC).

Bijleveld, M. (1974). The systematic persecution: A review of historical and more recent examples of the destruction of birds of prey in europe. In M. Bijleveld (Ed.), Birds of prey in europe (pp. 1–43). Macmillan Education, London. 10.1007/978-1-349-02393-6_1

Bloom, P., & Clark, W. (2001). Molt and sequence of plumage of golden eagles and a technique for in-hand ageing. North American Bird Bander, 3 (26), 97–116.

Bohrer, G., Brandes, D., Mandel, J. T., Bildstein, K. L., Miller, T. A., Lanzone, M., Katzner, T., Maisonneuve, C., & Tremblay, J. A. (2012). Estimating updraft velocity components over large spatial scales: Contrasting migration strategies of golden eagles and turkey vultures. Ecology Letters, 15 (2), 96–103. 10.1111/j.1461-0248.2011.01713.x

Brown, C. R., & Brown, M. B. (2013). Where has all the road kill gone? [Publisher: Elsevier]. Current Biology, 23 (6), R233–R234. 10.1016/j.cub.2013.02.023

Buskirk, J. V. (2012, June 14). Behavioural plasticity and environmental change. In U. Candolin & B. B. Wong (Eds.), Behavioural responses to a changing world: Mechanisms and consequences (p. 0). Oxford University Press. 10.1093/acprof:osobl/9780199602568.003.0011

Calenge, C. (2006). The package adehabitat for the r software: Tool for the analysis of space and habitat use by animals. Ecological Modelling, 197, 1035. 10.1016/j.ecolmodel.2006.03.017

Campbell-Staton, S. C., Arnold, B. J., Gonçalves, D., Granli, P., Poole, J., Long, R. A., & Pringle, R. M. (2021). Ivory poaching and the rapid evolution of tusklessness in african elephants [Publisher: American Association for the Advancement of Science]. Science, 374 (6566), 483–487. 10.1126/science.abe7389

Caspi, T., Johnson, J. R., Lambert, M. R., Schell, C. J., & Sih, A. (2022). Behavioral plasticity can facilitate evolution in urban environments. Trends in Ecology & Evolution, 37 (12), 1092–1103. 10.1016/j.tree.2022.08.002

Center, G. A. (2021). World settlement footprint (wsf) 2019 – sentinel-1/2 – global [10 m global settle-ment mask derived from Sentinel-1/2 imagery]. DLR. 10.15489/twg5xsnquw84

Chambert, T., Imberdis, L., Couloumy, C., Bonet, R., & Besnard, A. (2020). Density dependence in golden eagle aquila chrysaetos fecundity better explained by individual adjustment than territory heterogeneity. Ibis, 162 (4), 1312–1323. 10.1111/ibi.12826

Charrad, M., Ghazzali, N., Boiteau, V., & Niknafs, A. (2014). NbClust: An R Package for Determining the Relevant Number of Clusters in a Data Set. Journal of Statistical Software, 61 (6), 1–36. 10.18637/jss.v061.i06

Cheng, Y., Fiedler, W., Wikelski, M., & Flack, A. (2019). “closer-to-home” strategy benefits juvenile survival in a long-distance migratory bird. Ecology and Evolution, 9 (16), 8945–8952. 10.1002/ece3.5395

Conover, D. O., Munch, S. B., & Arnott, S. A. (2009). Reversal of evolutionary downsizing caused by selective harvest of large fish. Proceedings of the Royal Society B: Biological Sciences, 276 (1664), 2015–2020. 10.1098/rspb.2009.0003

Conrad, O., Bechtel, B., Bock, M., Dietrich, H., Fischer, E., Gerlitz, L., Wehberg, J., Wichmann, V., & Böhner, J. (2015). System for automated geoscientific analyses (saga) v. 2.1.4. Geoscientific Model Development, 8 (7), 1991–2007. 10.5194/gmd-8-1991-2015

Davis, J. M., & Stamps, J. A. (2004). The effect of natal experience on habitat preferences. Trends in Ecology & Evolution, 19 (8), 411–416. 10.1016/j.tree.2004.04.006

Donelan, S. C., Hellmann, J. K., Bell, A. M., Luttbeg, B., Orrock, J. L., Sheriff, M. J., & Sih, A. (2020). Transgenerational plasticity in human-altered environments. Trends in Ecology & Evolution, 35 (2), 115–124. 10.1016/j.tree.2019.09.003

Dugatkin, L. A. (2018). The silver fox domestication experiment. Evolution: Education and Outreach, 11 (1), 16. 10.1186/s12052-018-0090-x

Dugatkin, L. A. (2024). Anthropogenic evolution: Humans are changing more than just the environments species inhabit. we are changing the species themselves. Scientific American, 330 (6), 62. 10.1038/scientificamerican062024-7uPmh9cnc1QVsFBkH4XO0M

Fabbri, G., Biello, R., Gabrielli, M., Torres Vilaça, S., Sammarco, B., Fuselli, S., Santos, P., Ancona, L., Peretto, L., Padovani, G., Sollitto, M., Iannucci, A., Paule, L., Balestra, D., Gerdol, M., Ciofi, C., Ciucci, P., Mahan, C. G., Trucchi, E., … & Bertorelle, G. (2025). Coexisting with humans: Genomic and behavioral consequences in a small and isolated bear population. Molecular Biology and Evolution, 42 (12), msaf292. 10.1093/molbev/msaf292

Fasce, P., Fasce, L., Villers, A., Bergese, F., & Bretagnolle, V. (2011). Long-term breeding demography and density dependence in an increasing population of golden eagles Aquila chrysaetos. Ibis, 153 (3), 581–591. 10.1111/j.1474-919X.2011.01125.x

Fauchald, P., & Tveraa, T. (2003). Using first-passage time in the analysis of area-restricted search and habitat selection. Ecology, 84 (2), 282–288.

Faure, L. (2025a). Golden eagle landscape use. 10.17617/3.AB1BDX

Faure, L. (2025b). Landscape_use_golden_eagle.github repository. https://github.com/LouiseFau/Landscape_use_golden_eagle.git

Fernández-Gil, A., Lamas, J. A., Ansola, L. M., Román, J., de Gabriel Hernando, M., & Revilla, E. (2023). Population dynamics of recovering apex predators: Golden eagles in a mediterranean landscape. Journal of Zoology, 319 (2), 99–111. 10.1111/jzo.13026

Gazzelloni, S. (2014). Demographic changes in the alps, report on the state of the alps (No. 5). Permanent Secretariat of the Alpine Convention. Austria.

Ghalambor, C., Angeloni, L., & Carroll, S. (2010). Behavior as phenotypic plasticity. In Evolutionary behavioral ecology (DF Westneat & CW Fox, pp. 90–107). Oxford University Press.

Haller, H. (1996). Der steinadler in graubünden: Langfristige untersuchungen zur populationsökologie von Aquila chrysaetos im zentrum der alpen. Ala, Schweizerische Gesellschaft für Vogelkunde und Vogelschutz, 9, 186.

Harmon, E. A., & Pfennig, D. W. (2021). Evolutionary rescue via transgenerational plasticity: Evidence and implications for conservation. Evolution & Development, 23 (4), 292–307. 10.1111/ede.12373

Hemery, A., Mugnier-Lavorel, L., Itty, C., Duriez, O., & Besnard, A. (2023). Timing of departure from natal areas by golden eagles is not constrained by acquisition of flight skills. Journal of Avian Biology, 8 (7), e03111. 10.1111/jav.03111

Hendry, A. P., Farrugia, T. J., & Kinnison, M. T. (2008). Human influences on rates of phenotypic change in wild animal populations. Molecular Ecology, 17, 20–29. 10.1111/j.1365-294X.2007.03428.x

Isik, O. K., Hong, J., Petrunin, I., & Tsourdos, A. (2020). Integrity analysis for GPS-based navigation of UAVs in urban environment. Robotics, 9 (3), 66. 10.3390/robotics9030066

Jenny, D., Denis, S., Cruickshank, S. S., Tschumi, M., Hatzl, J., & Haller, H. (2024). The golden eagle in switzerland. In D. H. Ellis, J. Bautista & C. H. Ellis (Eds.), The golden eagle around the world: A monograph on a holarctic raptor (pp. 367–394). Hancock House Publishers, Surrey (BC).

Jolliffe, I. T. (2002). Principal component analysis (2nd edn). 271 pp. 10.1007/978-1-4757-1904-8

Karabanina, E., Lansink, G. M. J., Ponnikas, S., & Kvist, L. (2024). A renewed glance at the palearctic golden eagle: Genetic variation in space and time. Ecology and Evolution, 14 (3), e11109. 10.1002/ece3.11109

Kassambara, A. (2017a). Practical guide to cluster analysis in r: Unsupervised machine learning (Vol. 1). 187 pp.

Kassambara, A. (2017b). Practical guide to principal component methods in r: PCA, m (CA), FAMD, MFA, HCPC, factoextra (Vol. 2). 155 pp.

Kenward, R. E. (2000). A manual for wildlife radio tagging (2nd edn). 325 pp.

Kuijper, B., & Hoyle, R. B. (2015). When to rely on maternal effects and when on phenotypic plasticity? Evolution, 69 (4), 950–968. 10.1111/evo.12635

Lê, S., Josse, J., & Husson, F. (2008). FactoMineR: An r package for multivariate analysis. Journal of Statistical Software, 25, 1–18. 10.18637/jss.v025.i01

Lefebvre, L. (1995). The opening of milk bottles by birds: Evidence for accelerating learning rates, but against the wave-of-advance model of cultural transmission. Behavioural Processes, 34 (1), 43–53. 10.1016/0376-6357(94)00051-H

Loder, N. (2005). Point of no return. Conservation in Practice, 6 (3), 124–129. 10.1111/j.1526-4629.2005.tb00179.x

Lyly, M. S., Villers, A., Koivisto, E., Helle, P., Ollila, T., & Korpimäki, E. (2015). Avian top predator and the landscape of fear: Responses of mammalian mesopredators to risk imposed by the golden eagle. Ecology and Evolution, 5 (2), 503–514. 10.1002/ece3.1370

Mahalanobis, P. C. (1936). On the generalised distance in statistics. Proceedings of the National Institute of Sciences of India, 2 (1), 49–55.

Martínez-Abraín, A., Jiménez, J., & Oro, D. (2019). Pax romana: ‘refuge abandonment’ and spread of fearless behavior in a reconciling world. Animal Conservation, 22 (1), 3–13. 10.1111/acv.12429

Martínez-Abraín, A., Jiménez, J., & Ferrer, M. (2021). Changes from cliff-to tree-nesting in raptors: A response to lower human persecution? Journal of Raptor Research, 55 (1), 119–123. 10.3356/0892-1016-55.1.119

McGarigal, K., Stafford, S., & Cushman, S. (2000). Multivariate statistics for wildlife and ecology research. 283 pp.

Murgatroyd, M., Photopoulou, T., Underhill, L. G., Bouten, W., & Amar, A. (2018). Where eagles soar: Fine-resolution tracking reveals the spatiotemporal use of differential soaring modes in a large raptor. Ecology and Evolution, 8 (13), 6788–6799. 10.1002/ece3.4189

Nebel, C., Gamauf, A., Haring, E., Segelbacher, G., Väli, Ü., Villers, A., & Zachos, F. E. (2019). New insights into population structure of the european golden eagle (aquila chrysaetos) revealed by microsatellite analysis. Biological Journal of the Linnean Society, 128 (3), 611–631. 10.1093/biolinnean/blz130

Nielson, R. M., Murphy, R. K., Millsap, B. A., Howe, W. H., & Gardner, G. (2016). Modeling late-summer distribution of golden eagles aquila chrysaetos in the western united states. PLOS One, 11 (8), e0159271. 10.1371/journal.pone.0159271

Nourani, E., Faure, L., Brønnvik, H., Scacco, M., Bassi, E., Fiedler, W., Grüebler, M. U., Hatzl, J. S., Jenny, D., Roverselli, A., Sumasgutner, P., Tschumi, M., Wikelski, M., & Safi, K. (2024). Developmental stage shapes the realized energy landscape for a flight specialist. eLife, 13. 10.7554/eLife.98818.2

OpenStreetMap contributors. (2025). Openstreetmap [Open Data Commons Open Database License (ODbL)]. OpenStreetMap Foundation.

Orgeret, F., Kormann, U., Catitti, B., Witczak, S., van Bergen, V. S., Scherler, P., & Grüebler, M. U. (2024). Imprinted habitat selection varies across dispersal phases in a raptor species. Scientific Reports, 14, 26656. 10.1038/s41598-024-75815-1

Pedrini, P., & Sergio, F. (2001). Golden eagle aquila chrysaetos density and productivity in relation to land abandonment and forest expansion in the alps. Bird Study, 48 (2), 194–199. 10.1080/00063650109461218

Peres-Neto, P. R., Jackson, D. A., & Somers, K. M. (2005). How many principal components? stopping rules for determining the number of non-trivial axes revisited. Computational Statistics & Data Analysis, 49 (4), 974–997. 10.1016/j.csda.2004.06.015

Pigliucci, M. (2001). Phenotypic plasticity: Beyond nature and nurture. 356 pp.

Potvin, D. A., & Mulder, R. A. (2013). Immediate, independent adjustment of call pitch and amplitude in response to varying background noise by silvereyes (zosterops lateralis). Behavioral Ecology, 24 (6), 1363–1368. 10.1093/beheco/art075

Réale, D., Reader, S. M., Sol, D., McDougall, P. T., & Dingemanse, N. J. (2007). Integrating animal temperament within ecology and evolution. Biological Reviews, 82, 291–318. 10.1111/j.1469-185X.2007.00010.x

Ricau, B., & Decorde, V. (2009). L’Aigle royal: biologie histoire et conservation. Situation dans le Massif central. 522 pp.

Riley, S. J., DeGloria, S. D., & Elliot, R. (1999). Index that quantifies topographic heterogeneity. Intermountain Journal of Sciences, 5 (1), 23–27.

Schwarz, G. (1978). Estimating the dimension of a model. The Annals of Statistics, 6 (2), 461–464. 10.1214/aos/1176344136

Sergio, F., Pedrini, P., Rizzolli, F., & Marchesi, L. (2006). Adaptive range selection by golden eagles aquila chrysaetos in a changing landscape: A multiple modelling approach. Biological Conservation, 133 (1), 32–41. 10.1016/j.biocon.2006.05.015

Sih, A., Ferrari, M. C. O., & Harris, D. J. (2011). Evolution and behavioural responses to human-induced rapid environmental change. Evolutionary Applications, 4 (2), 367–387. 10.1111/j.1752-4571.2010.00166.x

Singh, N. J., Moss, E., Hipkiss, T., Ecke, F., Dettki, H., Sandström, P., Bloom, P., Kidd, J., Thomas, S., & Hörnfeldt, B. (2016). Habitat selection by adult golden eagles aquila chrysaetos during the breeding season and implications for wind farm establishment. Bird Study, 63 (2), 233–240. 10.1080/00063657.2016.1183110

Smith, B. R., & Blumstein, D. T. (2013). Chapter 13. animal personality and conservation biology: The importance of behavioral diversity. In C. Carere & D. Maestripieri (Eds.), Animal personalities: Behavior, physiology, and evolution (pp. 381–413). University of Chicago Press.

Tack, J. D., Noon, B. R., Bowen, Z. H., & Fedy, B. C. (2020). Ecosystem processes, land cover, climate, and human settlement shape dynamic distributions for golden eagle across the western us. Animal Conservation, 23 (1), 72–82. 10.1111/acv.12511

Thurfjell, H., Ciuti, S., & Boyce, M. S. (2017). Learning from the mistakes of others: How female elk (cervus elaphus) adjust behaviour with age to avoid hunters. PLOS ONE, 12 (6), 1–20. 10.1371/journal.pone.0178082

Tuomainen, U., & Candolin, U. (2011). Behavioural responses to human-induced environmental change. Biological Reviews, 86 (3), 640–657. 10.1111/j.1469-185X.2010.00164.x

Warnock, S., & Griffiths, G. (2015). Landscape characterisation: The living landscapes approach in the UK. Landscape Research, 40 (3), 261–278. 10.1080/01426397.2013.870541

Watson, J. (2010). The golden eagle (2nd edn). 439 pp.

Weiss, A. Topographic position and landforms analysis. In: In Esri user conference, vol? 200, san diego (ca). 2001.

Wentworth, A., Ricau, B., Itty, C., Bayle, P., Drillat, B., Couloumy, C., Clouet, M., & Seguin, J.-F. (2024). The golden eagle in france. In D. H. Ellis, J. Bautista & C. H. Ellis (Eds.), The golden eagle around the world: A monograph on a holarctic raptor (pp. 283–303). Hancock House Publishers.

Whitfield, D. P., Fielding, A. H., Gregory, M. J. P., Gordon, A. G., McLeod, D. R. A., & Haworth, P. F. (2007). Complex effects of habitat loss on golden eagles aquila chrysaetos. Ibis, 149 (1), 26–36. 10.1111/j.1474-919X.2006.00591.x

Wong, B. B. M., & Candolin, U. (2015). Behavioral responses to changing environments. Behavioral Ecology, 26 (3), 665–673. 10.1093/beheco/aru183

Zimmermann, S.-S., Grüebler, M. U., Hatzl, J. S., Bassi, E., Safi, K., Fiedler, W., Jenny, D., & Tschumi, M. (2025). Cascading carry-over effects of early activity phenotypes in golden eagles. bioRxiv. 10.1101/2025.09.21.677571

